# SpaceBio Knowledge Hub: A LiteratOmics Platform for Microgravity and Space Biology Research

**DOI:** 10.64898/2026.07.13.737239

**Authors:** Jose Cleydson F Silva, Arthur Vieira, Melinda S. Chue Donahey, Sirlane Maria do Carmo Silva, Tomas Veloso, Adilson Lopes, Nick Sexson, Richard Barker, D Marshall Porterfield, Carlos A. Silva, Raquel Dias

## Abstract

Space biology literature is growing exponentially. Existing infrastructure has not kept pace with organizing, synthesizing, and disseminating this knowledge. We present SpaceBio SpaceBio Knowledge Hub (www.spacebio.space), an integrated digital ecosystem that combines artificial intelligence, real-time data integration, and open-access infrastructure to advance research, education, and collaboration in microgravity, space biology and space exploration. The platform applies AI-driven approaches including natural language processing, machine learning, and automated content generation to construct a semantic atlas of the field. The atlas reveals the hierarchical thematic organization underlying microgravity-induced biological responses, space mission infrastructure, planetary science, and astrobiology. As part of this effort, SpaceBio is moving toward the construction of a LiteratOmics framework for microgravity, and space biology a systematic, AI-enabled approach to mining, integrating, and structuring the primary literature generated by omics-driven spaceflight research, treating the scientific literature itself as a navigable data layer alongside genomic, transcriptomic, and proteomic datasets. Built on a scalable, cloud-based architecture with a user-centered interface, SpaceBio supports literature exploration, data integration, and knowledge discovery for researchers, educators, students, industry partners, and citizen scientists. The platform also functions as a community-building ecosystem. It integrates hands-on research initiatives, AI-generated educational content, pilot data science projects, and social responsibility programs that broaden participation without compromising scientific rigor. AI-enabled digital environments can transform fragmented literature into a navigable knowledge landscape. SpaceBio accelerates research productivity, strengthens STEM education, and supports the global space life sciences community as human space exploration enters in the most ambitious era.

## Background and importance

Space exploration is not merely the future of humanity but represents the ultimate challenge that redefines our limits, fosters innovation, and propels us beyond the seemingly impossible. As missions advance toward long-term habitation on the Moon and Mars, reliance on Earth for water refueling becomes unsustainable, making in-situ resource utilization critical [^1,2,3,4^]. Miniaturized and autonomous systems are vital for microbiological research and water management aboard spacecraft [^5,6,7^]. The International Space Station (ISS) exemplifies humanity’s capability to operate in extreme environments, where microgravity, elevated CO₂ levels, and solar radiation exacerbate challenges such as microbial adaptation and astronaut health risks [^8^]. Research reveals that spaceflights can alter neuronal morphology and impair waste removal in the brain, necessitating strategies to safeguard the human nervous system [^9^]. Simultaneously, emerging technologies like microgravity-enabled fermentation hold promise for advancing research and supporting prolonged missions [^10^]. Efforts such as JAXA’s MMX mission to Mars’ moons, NASA’s Artemis program, China’s advancements in orbital stations, and India’s lunar missions highlight the progress of space engineering [^1,11,12,13,14^]. Nevertheless, with growing public interest in human space exploration, addressing these scientific and technological challenges through innovation and rigorous testing on off-Earth platforms is essential to ensure future mission safety and long-term viability.

China has also emerged as a major contributor to space exploration. The Tiangong Space Station became fully operational in 2022, supporting long-duration human missions and experiments in life sciences, materials science, and biotechnology^15,16,17^. The Chang’e lunar program achieved several milestones, including the first soft landing on the far side of the Moon (Chang’e-4) and two lunar sample-return missions (Chang’e-5 and Chang’e-6)^18^. The Tianwen program extended these capabilities into deep space. Tianwen-1 successfully deployed an orbiter, lander, and rover on Mars in a single mission^19^. The achievements have expanded opportunities for international scientific collaboration and generated valuable datasets for future lunar and interplanetary exploration.

Space exploration spans a wide array of disciplines, including science, technology, engineering, and mathematics (STEM), tackling challenges such as jet propulsion, the creation of bioregenerative life support systems (BLSS), and groundbreaking advancements in space biology [^20,21,22,23^] (https://eden-iss.net/). A significant amount of research has been conducted over the years to advance space exploration, particularly during the era of the Space Race, which began with the landmark launch of Sputnik 1 on October 4, 1957 [^24^]. This event marked the start of the Space Age and triggered rapid technological advancements, including sending humans to space and landing on the Moon [^25,26^]. The assembly of International Space Station (ISS) represents a major milestone in international collaboration, involving five space agencies: NASA (United States), Roscosmos (Russia), ESA (Europe), JAXA (Japan), and CSA (Canada). The ISS has enabled thousands of experiments across diverse fields such as space biology [^27,28,29^], engineering [^30^,], material science [^31^], and more [^32,33^]. Notable experiments include research on artificial retinas [^34,35^], biomining [^36^], and methods for growing plants in microgravity [^37,38^]. The scientific outcomes from ISS research have been extensive, with over 3,000 experiments contributing to advancements in technology and knowledge [^39,40,41,42,43^]. These results have been published in numerous scientific journals across various disciplines, furthering our understanding of microgravity’s effects and supporting future deep-space exploration missions.

Since then, an essential effort was applied to harbor computational platformer to extract, transform and standardize the data to become available to scientific community. The NASA Open Science Data Repository (OSDR) centralizes space-related biological and health data by integrating the Ames Life Sciences Data Archive and Gene Lab, an open-access platform for managing and analyzing multi-omics data like transcriptomics, proteomics, metabolomics, and epigenomics [^43^]. Guided by FAIR principles (findable, accessible, interoperable, reusable), OSDR facilitates global collaboration and bioinformatics queries. The Space Omics and Medical Atlas (SOMA) project aims to standardize biological measurements and facilitate the sharing of omics data from astronauts, including those participating in critical missions such as Inspiration4, Polaris Dawn, and Axiom [^42^]. Space exploration databases encompass diverse domains, each tailored to specific scientific objectives. For planetary missions, repositories such as the NASA Planetary Data System (PDS), ESA’s Planetary Science Archive (PSA), ISRO Science Data Archive, JAXA Data Archives, and Roscosmos’ IKI Science Data serve as pivotal sources of mission data. In astronomy and deep space observation, notable databases include SIMBAD, the NASA Exoplanet Archive, HEASARC (https://www.heasarc.gsfc.nasa.gov), the ALMA Science Archive [^44^], and SDSS, which specialize in cataloging celestial objects and astronomical datasets. Astrobiology and exobiology research benefit from resources like the NASA Astrobiology Institute and PDS Astrobiology, focusing on the study of life in the universe [^45^]. For satellite tracking and space missions, platforms such as CelesTrak, Space-Track, and the Minor Planet Center [^46^] monitor orbital objects and smaller celestial bodies. Finally, lunar and Martian exploration is supported by data from the Lunar Reconnaissance Orbiter, NASA Mars Science Laboratory (Curiosity Rover - https://www.science.nasa.gov/mission/msl-curiosity), and the USGS Astrogeology Science Center (https://www.usgs.gov/centers/astrogeology-science-center), which provide geological insights and mission-specific information [^47^].

Despite extensive efforts to organize and share space exploration data, a significant gap remains in the form of specialized, centralized databases dedicated to space exploration literature. Critical areas such as spaceflight, microgravity research, astrobiology, and exoplanet exploration lack unified repositories that could streamline information access and promote collaborative innovation. Establishing a Space Exploration Knowledge Hub could bridge this gap by serving as a comprehensive resource for scientists, engineers, technologists, mathematicians, citizen scientists, and enthusiasts. This knowledge hub would integrate diverse datasets and publications into a single platform, facilitating interdisciplinary collaboration and accelerating advancements in addressing the complex challenges of deep space missions and sustaining human life beyond Earth. Here, we present the SpaceBio Knowledge Hub (https://www.spacebio.space), a centralized platform designed to aggregate, organize, and disseminate critical research on space biosciences, fostering collaboration and accelerating discoveries in space health, astrobiology, and human adaptation to extraterrestrial environments. Featuring an interactive and user-friendly dashboard, the platform offers a responsive design with intuitive navigation and enhanced accessibility. It is designed to connect the STEM community, educators, and citizen scientists, creating an inclusive space for knowledge exchange and innovation. Beyond serving as a research repository, the SpaceBio Knowledge Hub actively promotes the development of data science projects, educational initiatives, and community engagement programs. The SpaceBio can facilitate interdisciplinary collaboration, it empowers users to contribute to groundbreaking research and inspires the next generation of space explorers.

## Methods

### System Architecture and Deployment Infrastructure of the SpaceBio Literature Database

The https://www.spacebio.space literature database backend modules were implemented using an MVC (Model-View-Controller) software architecture pattern [^48^], programmed in the python programming language using Django framework [^49^] and with PostgreSQL server relational databases [^50^]. The front-end implementation using Tailwind CSS (https://www.tailwindcss.com) in conjunction with Django framework templates, adhering to a modular and reusable design philosophy. This approach emphasized the componentization of user interfaces, to a cohesive and scalable design system. Tailwind’s utility-first classes significantly accelerated development, minimizing the need for custom CSS and streamlining code maintenance. The synergy between Tailwind and Django templates improved overall performance and developer productivity. We use Gunicorn as a WSGI server and Nginx as a reverse proxy to deploy the application effectively [^51^]. Gunicorn handles multiple connections by spawning worker processes, ensuring efficient request management. Nginx enhances this setup by serving as a reverse proxy, improving performance through load balancing, caching, and serving static files. It also strengthens security by providing SSL termination and shielding the application from direct client access. This combination ensures a secure, scalable, and production-ready deployment.

### Unified Data Collection Strategy: Literature, Launch Events, Media, and Sky Observations

The data warehouse structure was organized in SQL tables. The SpaceBio web crawler and data-driven were implemented using Europe PMC, PubMed [^52^], Scopus (https://www.scopus.com), Croosref (https://www.crossref.org) and Semantic Scholar (https://www.semanticscholar.org) APIs respectively. The search process employs carefully curated keywords to ensure high relevance and precision of the retrieved results. To optimize data collection while respecting usage policies, all API requests adhere to the standard rate limits imposed by each platform. This approach guarantees a thorough and efficient gathering of pertinent scientific information across diverse sources, providing a robust foundation for further analysis and research.

Comprehensive data on space missions and rocket launches is systematically sourced from two authoritative APIs: Spaceflight News and The Space Devs. These platforms provide real-time, detailed information on space events, ensuring up-to-date coverage. Key data points, including Mission details, Rocket specifications, Launch Site (Pad) information, and Involved Agencies, are individually extracted and meticulously organized into a structured database, facilitating efficient retrieval and analysis. Furthermore, the system incorporates associated YouTube links, enabling quick access to live streams and related video content for each launch. This integrated approach delivers a thorough and current overview of global space activities, enhancing user engagement and information accessibility.

The Open source Stellarium web was utilized to provide access to astronomy data [^53^]. The access to astronomy data, such as planets, moons, stars, constellations, and other celestial objects, is obtained using Stellarium Web. The tool provides detailed and real-time astronomical information, enabling precise observation, simulation, and study of the night sky.

### Large-Scale Semantic Analysis and Biological Knowledge Discovery Framework

We developed a scalable GPU-accelerated framework for large-scale semantic analysis of scientific literature, integrating transformer-based models, unsupervised clustering, manifold learning, and automated biological annotation to identify latent themes, characterize semantic structure, and generate interpretable labels from millions of documents. Documents were encoded into dense embeddings using a domain-specific transformer trained on SciBERT and S-BioBert biomedical corpora^54,55,56^. Representations were generated via attention-aware mean pooling and stored in memory-mapped arrays for efficient large-scale processing.

To improve cluster stability, outliers were removed based on distance from the global centroid using percentile filtering. Thresholds were optimized via internal metrics such as Silhouette^57^, Davies–Bouldin index^58^, Calinski–Harabasz^59^ combined into a composite index. Filtered embeddings were clustered using GPU-accelerated K-means^60^ (RAPIDS^61,62^/cuML), exploring multiple resolutions to capture thematic granularity. Cluster quality was assessed with cosine-based silhouette and complementary validation metrics. For visualization, embeddings were projected using UMAP^63^ into 2D and 3D spaces, preserving local and global relationships for intuitive exploration of scientific domains.

Biological interpretation used a semantic phrase-ranking framework: candidate concepts were extracted via entity recognition, noun-phrase mining, and domain filtering, embedded in the same space, and ranked by similarity to cluster centroids. Top phrases generated biologically meaningful cluster annotations. Hierarchical subclustering further resolved thematic heterogeneity, identifying specialized topics within broader domains. Additionally, k-nearest-neighbor^64^ semantic networks were constructed from embeddings to provide graph-based representations for analysis and visualization.

### Fine-Tuning Pipeline for GPT-2-Large: From Dataset Preparation to Optimized Model Training

The fine-tuning of the GPT-2 model was conducted using a structured pipeline to adapt the pre-trained architecture for domain-specific text generation tasks. Below, we detail the methodology in a format suitable for scientific reporting, emphasizing reproducibility and clarity.

The fine-tuning process utilized the *AutoModelForCausalLM* class from the hugging face transformers library, which is optimized for autoregressive language modeling. The GPT-2 tokenizer was initialized using the *AutoTokenizer* class, with modifications to address its lack of native padding functionality. Specifically, the *pad_token* was aligned with the *eos_token* to prevent tokenization errors during preprocessing. The vocabulary size of the tokenizer was updated to ensure compatibility with the pre-trained model’s parameters.

A custom text corpus was imported from a local .txt file using the Hugging Face Datasets library. The dataset was formatted into tokenized sequences with a maximum length of 128 tokens per example. Tokenization included truncation and padding operations applied in batches through the map function, ensuring uniform input dimensions required by autoregressive models.

Fine-tuning was executed using the Hugging Face Trainer API, which integrates multiple features for efficient model training. A *DataCollatorForLanguageModeling* was employed with *mlm=False,* maintaining GPT-2’s autoregressive paradigm. Training hyperparameters included: i) Epochs: 3; ii) batch size per device: 2 iii) Mixed precision optimization: enabled (*fp16=True*) for GPUs supporting half-precision floating-point operations; iv) checkpointing: Automatic saving every 1,000 steps, retaining only two checkpoints simultaneously to optimize storage usage.

Hyperparameter optimization was conducted using the Optuna framework, which utilizes an efficient search algorithm to explore the parameter space and identify the optimal configuration for fine-tuning the GPT-2-large model. The process began with the definition of an objective function that assessed the model’s performance based on perplexity (calculated as exp(eval_loss)), a metric particularly relevant for causal language models. The search space encompassed three key parameters: (1) per_device_train_batch_size, representing the training batch size, tested with discrete values of 2, 4, and 8; (2) learning_rate, sampled on a logarithmic scale between 5e-5 and 5e-4 to enhance sensitivity at lower magnitudes; and (3) num_train_epochs, specifying the number of training epochs, ranging from 1 to 5 to balance computational cost and performance. Each trial involved training the model with parameters proposed by Optuna, followed by evaluation on the validation set, where perplexity served as the optimization metric. A total of 10 search iterations were conducted, leveraging Optuna’s default Tree-structured Parzen Estimator (TPE) sampling strategy, which focuses on promising regions of the parameter space based on prior trials. Ultimately, the configuration yielding the lowest perplexity was selected for final training, ensuring a balance between computational efficiency and model quality. This systematic approach facilitated robust optimization by adapting to the nonlinearities inherent in transformer training and maximizing the generalization capabilities of the fine-tuned model.

The model was fine-tuned without intermediate evaluations or validation during training, ensuring a streamlined optimization process. Progress was monitored through logs generated at intervals of 500 steps. After the optimization phase, the best hyperparameters identified were applied in a final training session using the trainer configured with Optuna’s recommendations. The fine-tuned model and tokenizer were saved locally in the directory enabling their reuse in domain-specific text generation tasks. Additionally, FP16 (mixed precision) training was employed when a GPU was available, significantly accelerating the process without compromising model quality.

### Embedding-Based Semantic Search Architecture: Vector Generation, Storage, and Query Matching

Vector embeddings were generated using the snowflake-arctic-embed2 model pulled from ollama (https://ollama.com/library/snowflake-arctic-embed2). Abstract texts were extracted from a JSON file of entries. Special characters (non-ascii) were removed for the purposes of embedding. Ollama was used with a local endpoint to generate vector embeddings of length 1024. These vectors were inserted into a PostgresSQL database using pgvector for downstream analysis. Similarity comparisons are done by embedding the users query using the same process as above then doing a vector similarity search against the collection of vectors in the database. A user-defined top N number of results can be retrieved which should represent the abstracts which are most semantically like the user’s query, and thus most likely to have the desired content.

## Results

### An Integrated, Real-Time Database for Space Science Research, Education, and Exploration

This comprehensive and self-sustaining database that integrates heterogeneous sources of information researcher papers related to space exploration, aggregating over 382,600 scientific publications across key domains such as spaceflight, microgravity research, astrobiology, and exoplanet exploration (Supplementary table 1). The database serves as a critical resource for the scientific community, educators and space enthusiasts, providing centralized access to contextualized data through an advanced search engine. Access to the database enables the extraction of detailed data regarding, for instance, plant development under microgravity conditions such as alterations in growth orientation, hormone distribution, and gene expression in response to the absence of gravity as well as insights into muscle physiology within extraterrestrial environments, including bone mass loss, muscle atrophy, and changes in musculoskeletal structure experienced by astronauts during prolonged space missions. Additionally, the database supports a wide range of STEAM (Science, Technology, Engineering, Arts, and Mathematics) research by providing rapid access to large, multidimensional datasets, thereby facilitating interdisciplinary studies in areas like biophysics, fluid dynamics, materials science, and beyond. The resource facilitates interdisciplinary collaboration and accelerates the transfer of scientific knowledge across related domains.

To ensure real-time relevance, the platform dynamically aggregates mission-specific data, including rocket launch schedules, payload details, experimental protocols, launch site information, and data on participating space agencies. This enables researchers to monitor ongoing or scheduled missions, promoting collaboration and alignment with current exploratory efforts. Additionally, the database curate’s news on spaceflight from reputable journalistic sources, offering professionals, academics, and enthusiasts a consolidated and continuously updated information hub (Figure 1).

**Figure 1.**
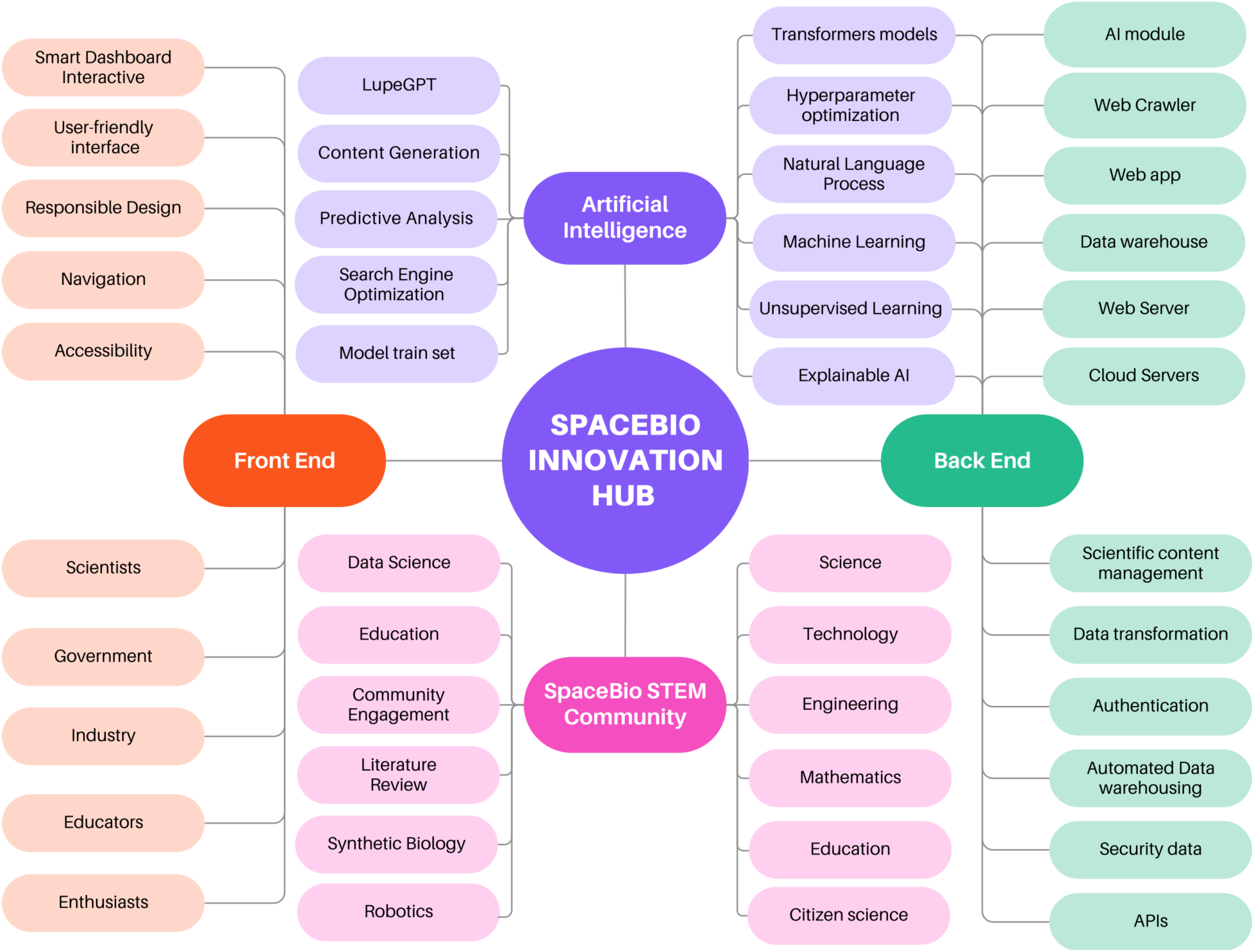
Architecture of the SpaceBio Innovation Hub. The platform integrates Artificial Intelligence, Front End, Back End, and the SpaceBio STEM Community. AI supports data analysis, content generation, and decision-making. The Front End provides an accessible user experience. The Back End manages data, security, cloud services, and APIs. The STEM community connects researchers, educators, industry, government, and citizens to foster collaboration and innovation in space biology and related fields.

A distinctive feature of the platform is SpaceBio TV, an integrated multimedia resource that archives historical space launch recordings via YouTube, thereby broadening educational outreach and accessibility. The platform also incorporates a virtual telescope tool powered by Stellarium Web, which provides an interactive educational interface for real-time celestial observation. Thisfunctionality allows educators and researchers to visualize satellites, planets, moons, and galaxies, bridging the gap between theoretical research and public engagement in astronomy.

Unifying fragmented data streams into a single searchable platform, the system not only enhances research efficiency but also democratizes access to space science, promoting global collaboration and scientific literacy. Future developments will include the application of machine learning to improve data retrieval and predictive analytics, further consolidating the platform’s role as an indispensable tool for the space science community.

### AI-enhanced Knowledge Discovery in Microgravity and Spaceflights Research

#### AI-Derived Semantic Atlas of Space Exploration and Space Biology

Space Exploration and Space Biology produce a rapidly expanding and highly interdisciplinary body of knowledge spanning biological sciences, medicine, engineering, planetary sciences, and aerospace technologies. To systematically investigate this domain, we developed an AI-driven framework that integrates transformer-based semantic embeddings and unsupervised learning to uncover latent conceptual relationships directly from approximately 0.5 million curated records. The resulting semantic landscape provides a quantitative representation of the structure of space-related research, enabling the identification of major scientific domains, interdisciplinary connections, and emerging research frontiers. We performed unsupervised learning analysis across the SpaceBio knowledge landscape systematically evaluated over k=10-50 using four complementary metrics: Silhouette score, Calinski-Harabasz index, Davies-Bouldin index, and a composite decision index integrating normalized scores. Increasing k produced progressively finer segmentation of the semantic space, accompanied by divergent metric behavior. The composite decision framework identified k=30 as the optimal clustering resolution, yielding the maximal decision index across all configurations (Figure 2). At this scale, the semantic partition achieved an optimal trade-off between intracluster cohesion, intercluster separation, and interpretability. While the Davies-Bouldin index continued to decrease at higher k, both Silhouette and Calinski-Harabasz metrics declined beyond K=30, indicating over-segmentation and reduced structural coherence. The convergence of independent evaluation criteria at k=30 supports a robust partitioning of the SpaceBio knowledge base into thirty discrete semantic domains. This resolution was therefore adopted for all subsequent analyses, including semantic annotation, hierarchical decomposition, and domain-specific interpretation of the knowledge landscape.

**Figure 2.**
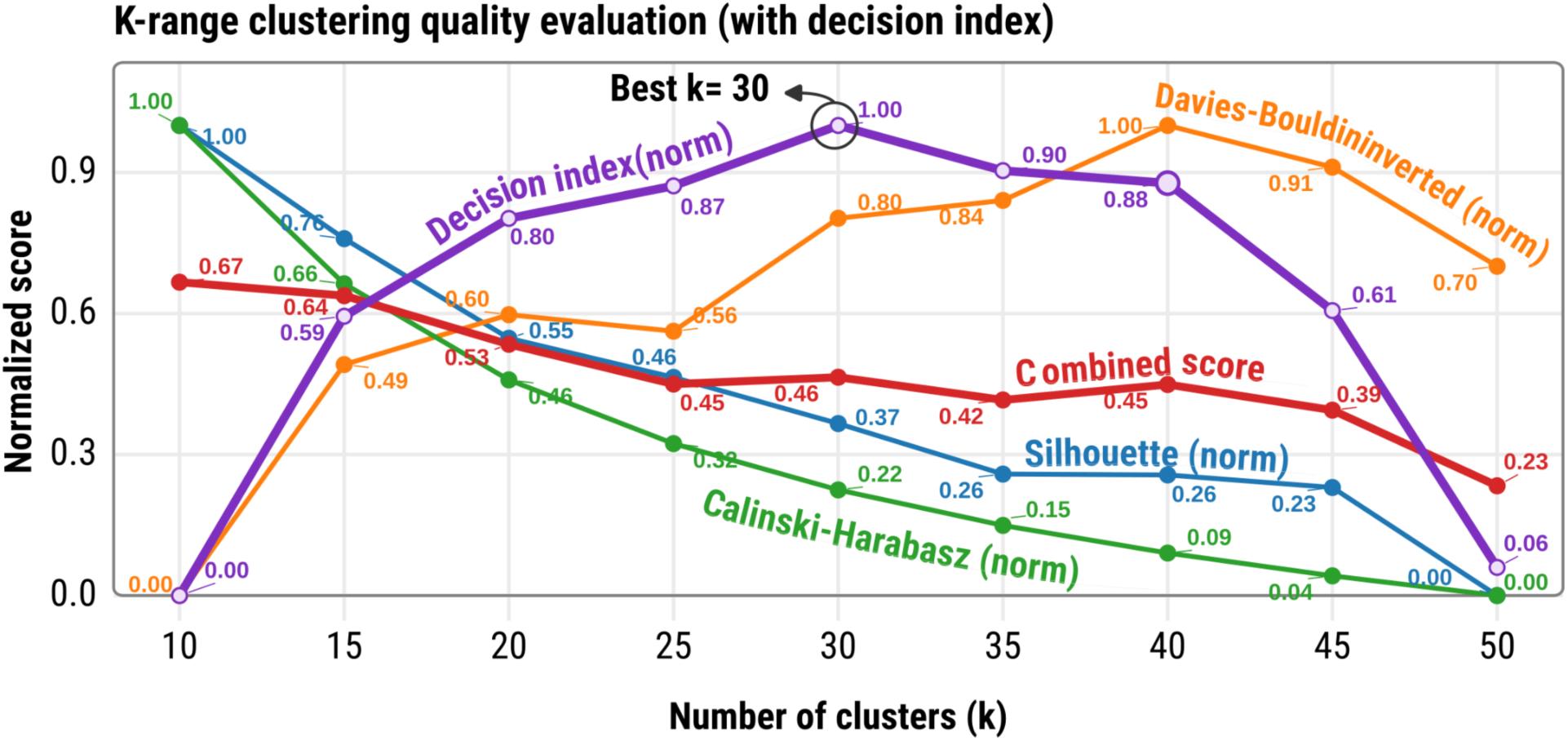
AI-guided optimization of semantic clustering across the SpaceBio knowledge landscape. Clustering performance was evaluated for k=10–50 using normalized Silhouette, Calinski-Harabasz, and inverted Davies-Bouldin indices, along with a composite decision index. The framework identified k=30 as the optimal resolution, achieving the best balance between cohesion, separation, and interpretability.

We applied an AI-driven framework to the SpaceBio knowledge base, combining semantic embeddings, dimensionality reduction, and clustering (Figure S1). This approach revealed a structured network spanning Space Exploration and Space Biology. We identified thirty major knowledge domains, including aerospace engineering, planetary sciences, materials research, and biological systems (Figure 2). The domains formed distinct interconnected clusters, reflecting the multidisciplinary nature of space research. Space Biology emerged as a central and highly connected domain. Biological processes such as microgravity adaptation, radiation response, and gene expression were strongly linked to technological and environmental fields, indicating deep integration across disciplines (Figure 3 - Cluster C11, C15, and C27). Microgravity acted as a key organizing principle. It connected research in biology, materials science, fluid physics, and engineering, highlighting its unifying role in long-duration spaceflight studies. All structures were derived directly from AI-learned semantic relationships, without manual classification. This demonstrates that large-scale language models can uncover the latent organization of scientific knowledge. Overall, the semantic landscape provides a quantitative atlas of Space Exploration and Space Biology and supports systematic knowledge discovery and interdisciplinary innovation.

**Figure 3.**
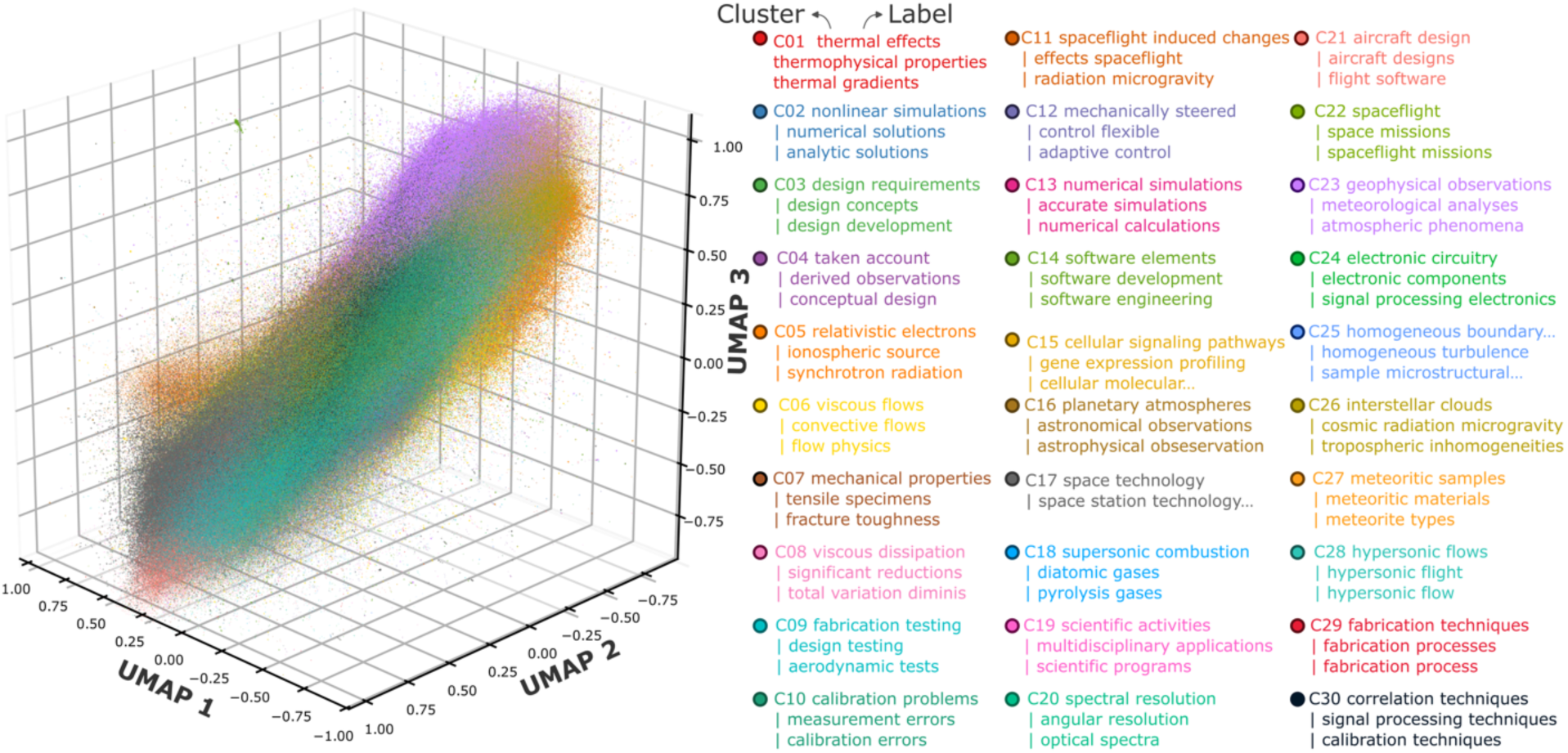
AI-generated semantic landscape of Space Exploration and Space Biology. Thirty knowledge domains identified using transformer-based embeddings and unsupervised clustering. The map shows interconnected regions across engineering, planetary sciences, space technologies, computational sciences, and biological systems, with microgravity and spaceflight adaptation at the core of the network.

#### Hierarchical Organization of Microgravity-Induced Biological Responses

To resolve the internal structure of spaceflight-induced biological responses, cluster C11 (Figure 4) was extracted from the global semantic landscape and reanalyzed using the same AI-driven framework. The cluster captures the intersection of microgravity, radiation, and physiological adaptation. High-resolution clustering partitioned C11 into 15 distinct subdomains, revealing a hierarchical organization not evident at the global level. The subdomains correspond to specialized research areas, including microgravity experiments, radiation effects, cardiovascular and musculoskeletal adaptation, plant biology, microbial responses, and long-duration spaceflight stressors (Figure 5). Microgravity-centered subdomains were prominent, underscoring its role as a primary driver of biological change across scales. Additional clusters captured molecular and systemic responses to radiation, as well as adaptive processes in plants and microorganisms under spaceflight conditions. This hierarchical structure indicates that spaceflight-induced biological responses comprise multiple interconnected domains. More broadly, it demonstrates that AI-based semantic analysis can resolve fine-grained organization within complex scientific fields, enabling the identification of specialized domains within Space Biology.

**Figure 4.**
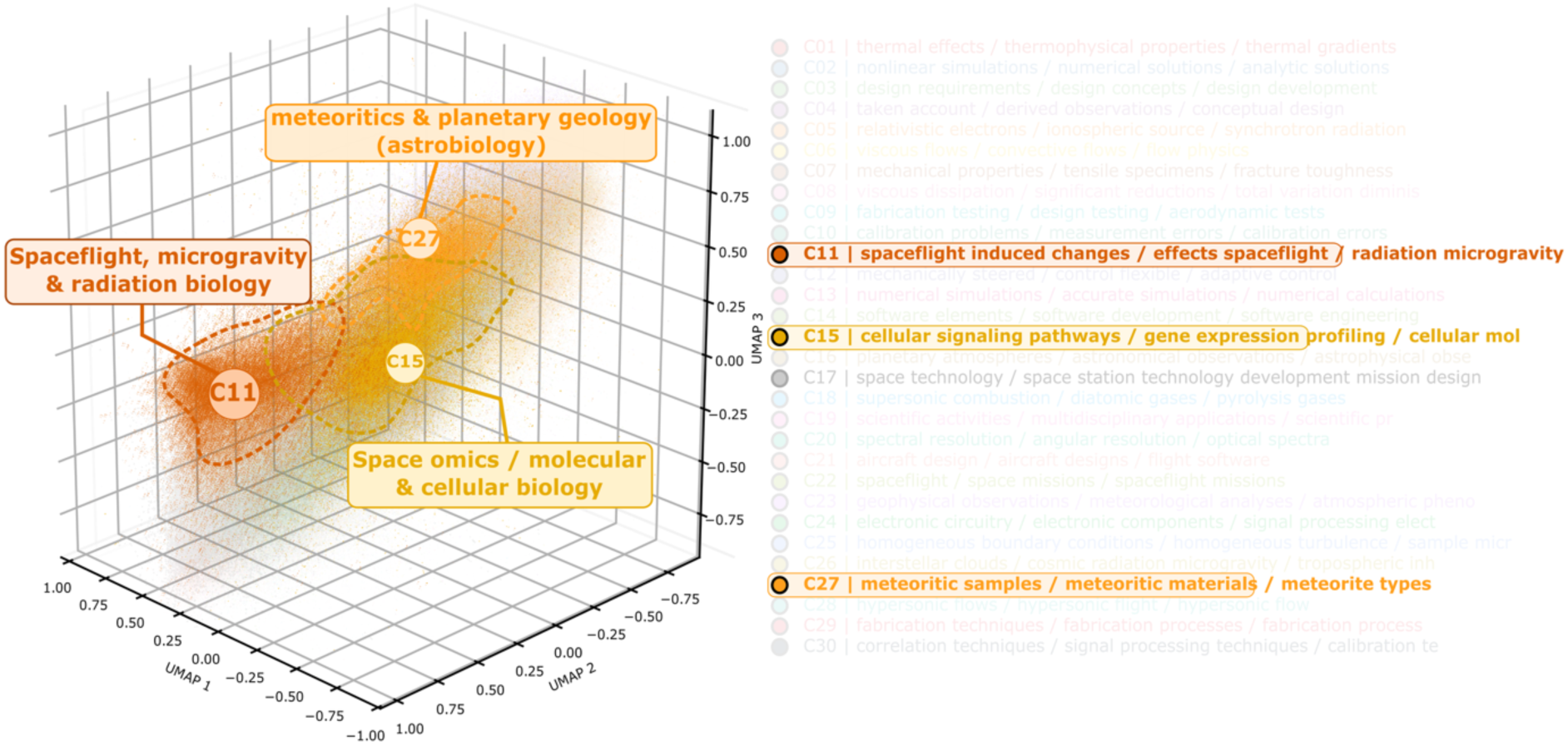
AI-driven semantic organization of biological domains within the SpaceBio landscape. Three-dimensional embeddings reveal relationships among key biological domains in space research. A prominent cluster of spaceflight-induced responses (C11), including microgravity adaptation and radiation processes, is closely linked to a molecular biology domain (C15) defined by cellular signaling and gene expression. In contrast, an astrobiology-related region (C27), focused on meteoritics and planetary geology, remains more distinct. The proximity of C11 and C15 highlights strong links between spaceflight stressors and molecular responses, while the separation of C27 reflects its specialized focus. These patterns show how AI-based analysis captures the structure and connectivity of Space Biology across the space science ecosystem.

**Figure 5.**
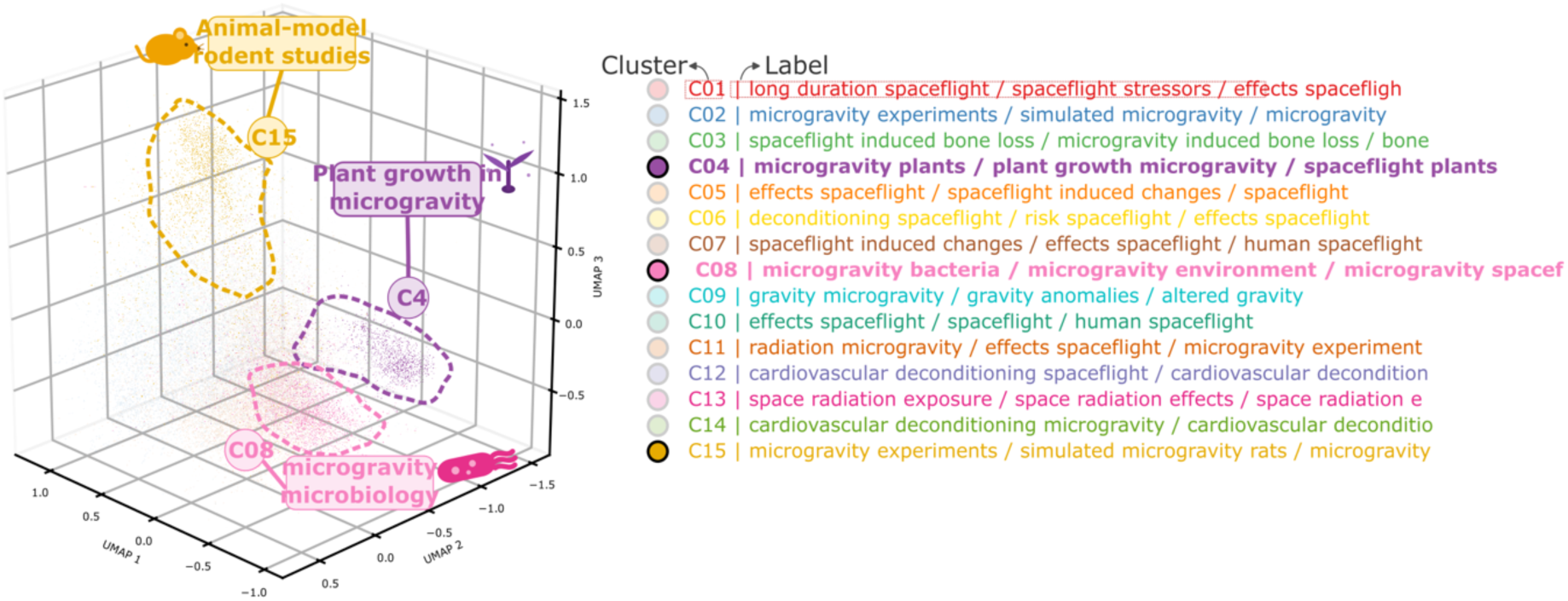
AI exposes the hidden architecture of biological adaptation to spaceflight. Recursive semantic analysis of the spaceflight response domain uncovered fifteen interconnected research communities spanning microgravity, radiation exposure, human physiology, plant biology, microbiology, and long-duration mission stressors.

Higher-order clustering identified four superordinate thematic domains (C15, C22, C26, and C27), each encompassing functionally coherent subclusters reflecting distinct areas of space biology and space research (Figure 6). Cluster C15 emerged as the primary biologically focused supercluster, integrating gene expression profiling and radiation-responsive transcriptomics, molecular radiation response under simulated microgravity, and vestibular and ocular changes, alongside cardiovascular, microbial, and musculoskeletal subclusters. Their tight spatial proximity indicated strong conceptual overlap between radiation biology, sensorimotor adaptation, and transcriptomic responses, suggesting these research lines are functionally intertwined within the broader spaceflight-induced biological adaptation literature.

**Figure 6.**
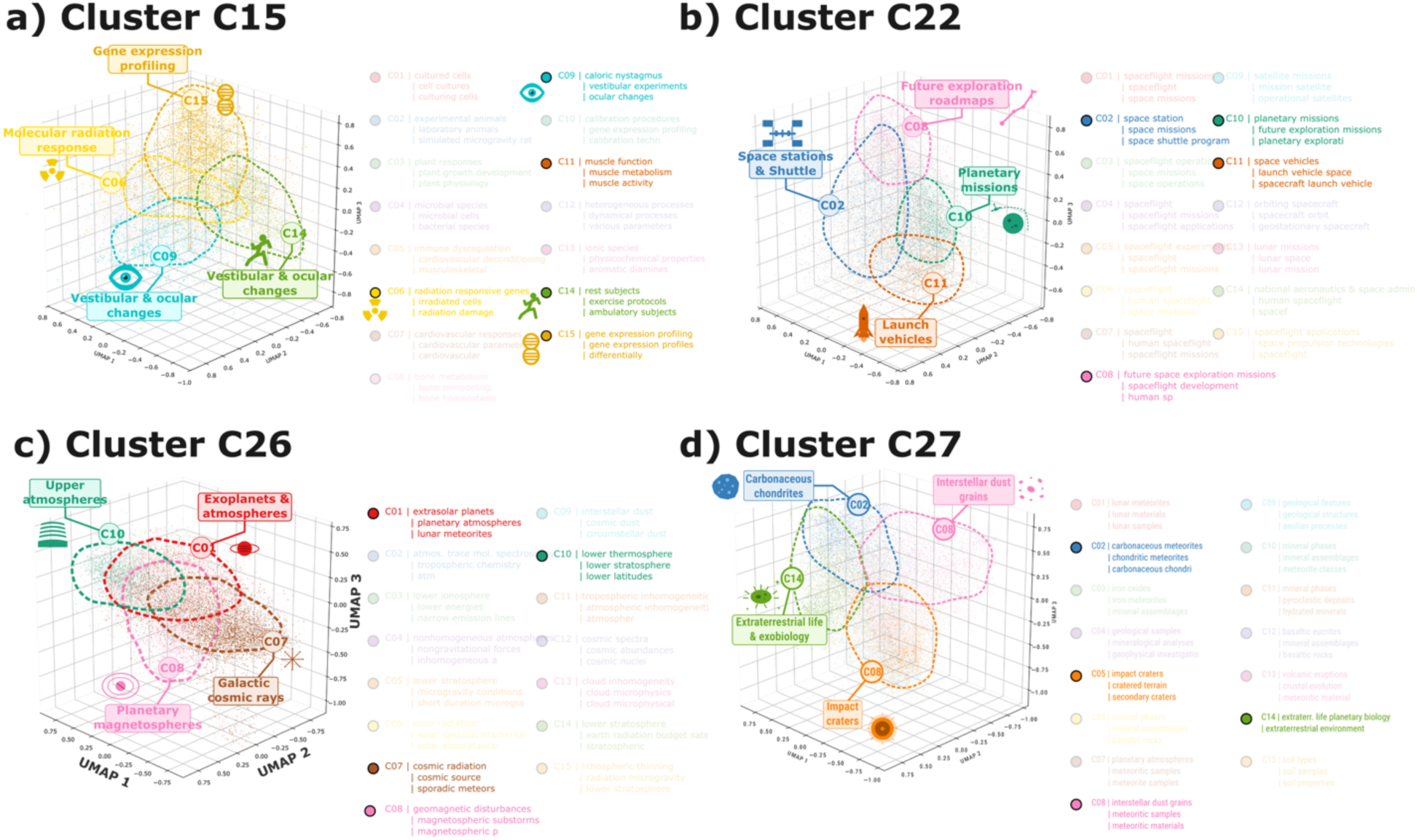
Hierarchical subclustering of major SpaceBio semantic domains. a) The molecular and cellular biology domain resolves into gene expression profiling, molecular radiation response, and vestibular and ocular changes. b) The space mission’s domain resolves into space stations and shuttle operations, launch vehicles, planetary missions, and future exploration roadmaps. c) The planetary atmospheres and astrophysics domain resolves into exoplanets and atmospheres, upper atmospheres, galactic cosmic rays, and planetary magnetospheres. d) The astrobiology and meteoritics domain resolves into carbonaceous chondrites, interstellar dust grains, impact craters, and extraterrestrial life and exobiology.

Higher-order clustering reveals the hierarchical thematic organization of the research landscape. Three-dimensional UMAP projections illustrate the macroorganization of the scientific corpus into four superordinate thematic domains, each encompassing functionally coherent subclusters reflecting conceptually distinct research orientations: biological and biomedical responses to microgravity, integrating transcriptomic adaptation, radiation-induced cellular damage, and sensorimotor alterations (C15; a); operational and technological dimensions of human spaceflight, spanning strategic mission planning and implementation (C22; b); physical and environmental contexts of spaceflight, from planetary atmospheric dynamics to deep-space radiation environments (C26; c); and planetary science, extraterrestrial materials, and astrobiology, where the spatial proximity of the exobiology subcluster to carbonaceous chondrite and interstellar dust grain groupings reflects conceptual linkages between prebiotic chemistry and primitive solar system materials (C27; d). Each point represents an individual document colored by subcluster identity (C01–C15), with dashed ellipses delimiting thematic groupings and lateral legends indicating the most representative terms of each subcluster.

Cluster C22 organized the operational and technological dimensions of human spaceflight into space station and shuttle operations, future exploration roadmaps, planetary mission objectives, and launch vehicle engineering. The spatial separation between strategic planning and mission-implementation subclusters reflected their conceptual distinction, while their co-membership within C22 underscored a shared organizational context within space exploration infrastructure. Cluster C26 captured planetary atmospheric physics, cosmic radiation, and magnetospheric phenomena, spanning exoplanetary atmospheres, upper atmospheric dynamics, galactic cosmic ray sources, and magnetospheric particle dynamics, unified by their relevance to the physical environments encountered during spaceflight.

Cluster C27 consolidated planetary science, extraterrestrial materials, and astrobiology into four thematic groupings: carbonaceous chondrite composition, interstellar dust grain and meteoritic sample analyses, impact crater morphology and secondary crater formation, and extraterrestrial life and exobiology. The positioning of the exobiology subcluster in proximity to the carbonaceous chondrite and interstellar dust grain groupings is consistent with conceptual linkages between primitive solar system materials and prebiotic chemistry, reinforcing the thematic coherence of this astrobiology-focused supercluster.

### Space Biology Outreach, Education, and Community Building

SpaceBio expanded beyond knowledge discovery to operate as an integrated scientific ecosystem supporting participation in Space Biology and Space Exploration. By combining artificial intelligence, open-access resources, collaborative research, and educational initiatives, the platform connected scientific discovery, workforce development, and public engagement across interdisciplinary communities encompassing students, educators, researchers, and citizen scientists working at the interface of microgravity biology, astrobiology, planetary habitability, and space biotechnology.

The platform supported hands-on research initiatives that created accessible entry points into Space Biology. Motivated by the cutting edge of space exploration particularly NASA’s Artemis III mission these initiatives included studies of microbial growth in modified Winogradsky columns using lunar and Martian regolith, examining how microorganisms colonize extraterrestrial soil under space-relevant conditions (Figure 7). Pilot projects extended the platform’s scientific scope in complementary directions (https://www.spacebio.space/spacebio_lab). A data science initiative developed an interactive dashboard for analyzing trends and identifying emerging opportunities in human space exploration, enabling data-driven visualization of research gaps and mission objectives. A semantic enrichment project applied Knowledge Graph methodologies and FAIR principles Findable, Accessible, Interoperable, and Reusable to enhance data discovery and interoperability within the SpaceBio platform, promoting advanced knowledge dissemination and fostering innovation across the research community. An educational initiative developed in partnership with SpaceBio engaged a doctoral candidate in a qualifying examination structured in a preprint draft format, exploring reverse ecology and microorganism prospecting for plant cultivation in space environments, fostering a practical connection between advanced research and training. Additionally, a literature review on the impacts of magnetic fields on living organisms examined perspectives on biomagnetism, broadening the platform’s engagement with integrative biological research relevant to spaceflight health challenges.

**Figure 7.**
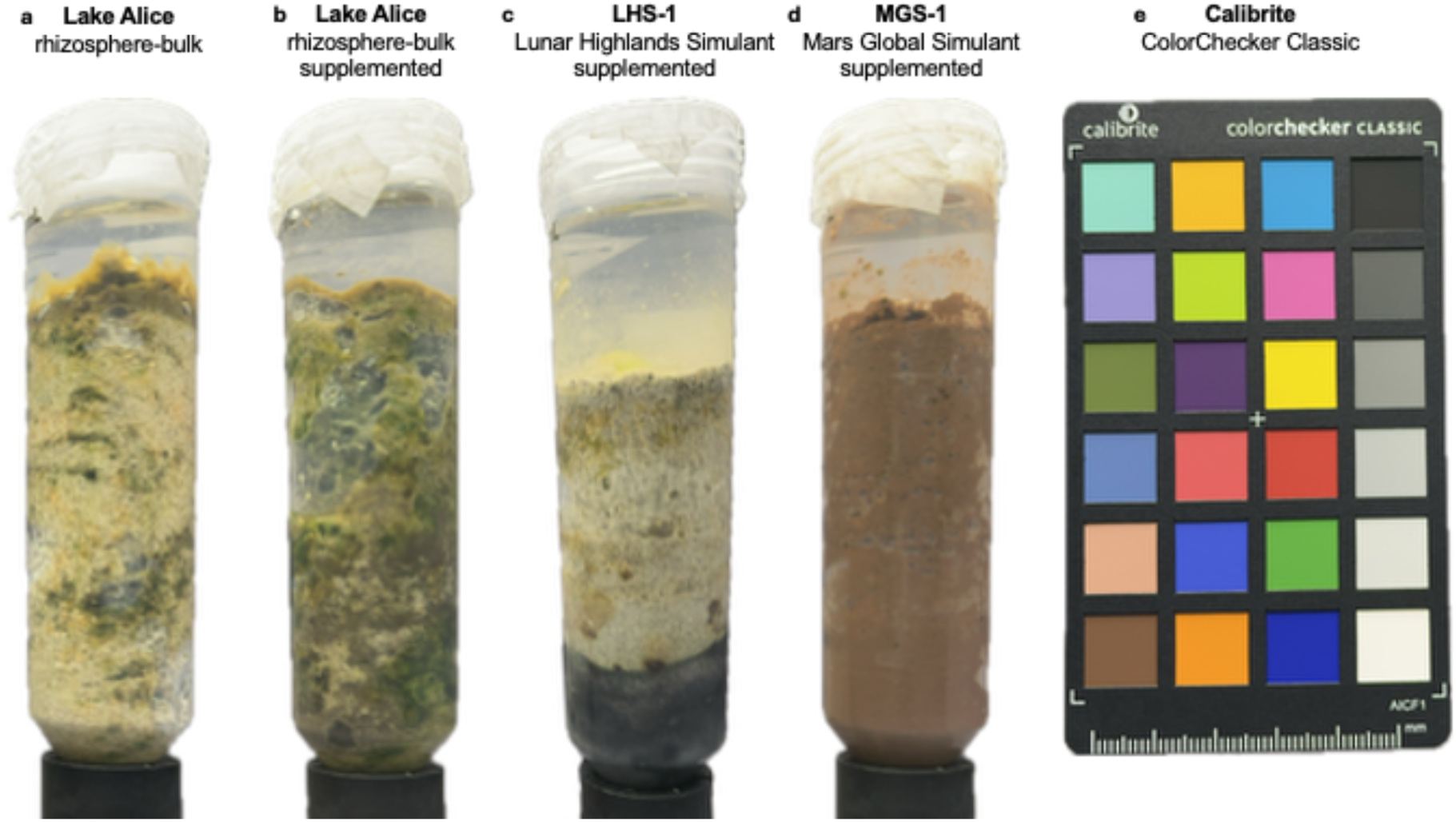
Figure 1. Ecophysiological stratification of Winogradsky columns inoculated with Lake Alice rhizosphere-bulk consortia and incubated in native sediment or extraterrestrial regolith simulants. (a) Lake Alice rhizosphere-bulk, unsupplemented. (b) Lake Alice rhizosphere-bulk, supplemented with starch and MgSO₄·7H₂O. Dense green phototrophic biofilm dominates the upper column, with a darker olive transition zone and scattered black patches at depth, indicating coexistence of phototrophic and sulfate-reducing guilds. (c) LHS-1 (Lunar Highlands Simulant), supplemented. Discrete vertical banding with a translucent beige-yellow aerobic surface zone overlying a sharply defined black basal layer, consistent with dissimilatory sulfate reduction and iron sulfide precipitation; phototrophic colonization is visibly suppressed relative to (b). (d) MGS-1 (Mars Global Simulant), supplemented. Homogeneous reddish-brown coloration throughout the column with no visible green zonation, consistent with dominance of iron-cycling metabolism (iron oxidation/reduction) and strong suppression of phototrophic growth. (e) Calibrite ColorChecker Classic, included for color calibration and standardization across image acquisitions.

Scientific communication constituted a major component of the platform, with artificial intelligence used to transform complex findings into educational content, visual narratives, and multimedia resources distributed through the SpaceBio digital ecosystem and social media channels, extending the reach of Space Biology beyond traditional academic audiences. The platform further supported international community formation through workshops, seminars, collaborative meetings, and conference participation, facilitating interaction among students, early-career researchers, and established investigators within the global Space Biology community. Consistent with its commitment to social responsibility and community engagement, SpaceBio implemented initiatives aimed at bridging the digital divide, including the donation of computers to underserved individuals to provide access to the platform and its innovation hub.

## Discussion

The rapid expansion of human spaceflight programs, including long-duration missions to the Moon and Mars, has intensified the need for systematic approaches to organizing and synthesizing the growing body of space biology knowledge^65^. The biological consequences of microgravity exposure are broad and molecularly complex, encompassing transcriptomic reprogramming, epigenetic modifications, proteomic shifts, and metabolic dysregulation that collectively challenge the homeostatic capacity of living systems in ways that remain incompletely understood^66^. Despite significant advances in omics technologies applied to spaceflight experiments from RNA sequencing of microgravity-exposed cell cultures to multi-omics profiling of astronaut cohorts the resulting data landscape remains fragmented across siloed, static repositories that were not designed to support integrative warehouse for harbor the literaturOmics across biological scales or disciplinary boundaries. SpaceBio was conceived as a structural response to this limitation, providing an integrated, real-time platform anchored in FAIR principles that unifies heterogeneous data sources spanning peer-reviewed literature, mission reports, and experimental records within a single openly accessible infrastructure capable of supporting both specialist research and broader scientific engagement.

Artificial intelligence was applied to the corpus organized within SpaceBio, natural language processing and unsupervised clustering transformed a vast and unstructured literature into a navigable semantic atlas an epistemological instrument that rendered the hidden intellectual architecture of space biology legible at scale. The resulting thematic map revealed that microgravity research is not a monolithic field but a structured landscape of functionally coherent domains, with gene expression profiling, radiation-responsive transcriptomics, and sensorimotor biology emerging as conceptually proximate research lines despite their historical pursuit in relative disciplinary isolation. The semantic organization has direct practical implications: by making the intellectual structure of the field visible, AI-driven analysis enables the identification of underexplored intersections between omics science and operational spaceflight medicine, highlights research areas where data generation has outpaced mechanistic interpretation and creates the conditions for hypothesis-driven investigation that transcends the boundaries of any single discipline.

The hierarchical clustering of microgravity-induced biological responses further demonstrated that the convergence of radiation biology, transcriptomics, and sensorimotor adaptation within a single supercluster is not a statistical artifact but a reflection of genuine molecular and physiological interdependencies. Ionizing radiation and microgravity act as co-stressors on the same cellular machinery inducing DNA damage responses, oxidative stress pathways, and inflammatory signaling cascades that intersect with the musculoskeletal, cardiovascular, and neurovestibular adaptations documented across astronaut omics studies^67,68^. Co-clustering argues for a systems biology framework in which microgravity-induced gene expression changes are interpreted not in isolation but in the context of the full environmental stressor profile of spaceflight, with implications for the development of multi-target countermeasures for long-duration missions such as Artemis^69,70^ and Mars expeditions^71,72,73^. The structural separation of planetary science, astrobiology, and space mission infrastructure into distinct superclusters further delineates where molecular biology ends and where geochemistry, engineering, and exploration strategy begin boundaries that are conceptually important for allocating research resources and designing interdisciplinary collaboration frameworks.

The biological complexity revealed by AI-driven analysis underscores that advancing space biology requires not only more data but better-connected communities capable of interpreting and acting upon it. SpaceBio therefore integrated outreach, education, and social responsibility as core functional dimensions of the platform, recognizing that the pipeline of researchers trained in microgravity biology, omics data science, and space medicine directly determines the field’s capacity to translate molecular findings into operational countermeasures. Pilot projects in semantic knowledge graph development, interactive data dashboards for space exploration trends, and educational initiatives connecting doctoral training to space biology applications collectively demonstrated that broadening scientific participation accelerates rather than dilutes research quality. As the molecular consequences of long-duration spaceflight become increasingly central to mission planning, the infrastructure required to synthesize, disseminate, and act upon that knowledge must grow in parallel making platforms such as SpaceBio not ancillary tools but foundational elements of the scientific enterprise that will carry human life beyond Earth.

### Challenges and Opportunities

A major challenge in space biology and microgravity research is the fragmentation of scientific knowledge across diverse repositories, databases, journals, and space agency platforms. Although initiatives such as NASA’s OSDR and GeneLab^74^ have improved data accessibility, critical information remains dispersed, limiting efficient discovery, integration, and reuse. The rapid growth of publications in areas such as space medicine, astrobiology, biotechnology, microbiology, and human spaceflight further complicates the identification of relevant findings and emerging trends. In addition, inconsistencies in metadata standards, data formats, and experimental protocols hinder interoperability and reproducibility, reducing the potential for large-scale analyses and cross-mission comparisons. Addressing these challenges requires innovative approaches that not only organize and connect scientific information, but also transform fragmented data into accessible, actionable, and collaborative knowledge.

The convergence of artificial intelligence, open science, and data-driven research offers a unique opportunity to overcome these limitations. The SpaceBio Knowledge Hub introduces the concept of LiteratOmics, a framework that integrates literature mining, natural language processing, knowledge graphs, and intelligent analytics to support scientific discovery. To unify publications, datasets, educational resources, and collaborative initiatives within a cohesive ecosystem, SpaceBio enhances knowledge accessibility and fosters interdisciplinary collaboration among researchers, educators, industry stakeholders, and citizen scientists. Beyond knowledge integration, the platform promotes the development of regional leadership networks and local communities worldwide, creating spaces for scientific dialogue, training, mentorship, and collaborative innovation. These distributed hubs can strengthen STEM engagement, support citizen science initiatives, and expand participation in space-related research across diverse geographic and socioeconomic contexts. As programs such as Artemis, Gateway, Tiangong, and future lunar and Martian missions generate unprecedented volumes of scientific data, platforms like SpaceBio will play a critical role in transforming information into insight, accelerating discoveries in microgravity and space biology, and empowering a globally connected community dedicated to the future of space exploration.

## Conclusion and future work

The SpaceBio Knowledge Hub is an AI-driven ecosystem integrating scientific literature, datasets, and educational resources to advance research in microgravity, space biology, and space exploration. Powered by the LiteratOmics framework, the platform accelerates knowledge discovery, fosters interdisciplinary collaboration, and connects scientific communities worldwide. Future efforts will expand data integration, advance AI capabilities, and strengthen regional leadership networks driving innovation, promoting STEM engagement, and enabling broader international collaboration in the space sciences.

## Acknowledgments

The authors gratefully acknowledge the contributions of all students, collaborators, and institutional partners whose engagement was essential to the realization of these initiatives, including the recipient communities and individuals whose participation reinforced SpaceBio’s commitment to inclusive and socially responsible science.

## Data Availability

The SpaceBio source code is publicly available at: https://github.com/jcleydsonsilva/Spacebio. A lightweight version of the platform is available for developers, educators, and researchers interested in exploring the framework or developing new applications and extensions based on the SpaceBio ecosystem.

## References

(1) Chatzitheodoridis, E.; De Vera, J.-P.; Kereszturi, A.; Mason, N.; Possnig, C.; Puumala, M.; Sivula, O.; Viso, M.; Arnould, J.; Detrell, G.; Ewald, R.; Freissinet, C.; Holynska, M.; Javaux, E.; Lee, N. M.; Lehto, K.; Milligan, T.; Siljeström, S.; Vago, J. L. Perspectives for Crewed Missions to Mars: Exploration from Orbit and/or Short Stay. In Mars and the Earthlings: A Realistic View on Mars Exploration and Settlement; Verseux, C., Gargaud, M., Lehto, K., Viso, M., Eds.; Space and Society; Springer Nature Switzerland: Cham, 2024; pp 117–197. 10.1007/978-3-031-66881-4_6.

(2) Kessler, P.; Prater, T.; Nickens, T.; Harris, D. Artemis Deep Space Habitation: Enabling a Sustained Human Presence on the Moon and Beyond. In 2022 IEEE Aerospace Conference (AERO); IEEE: Big Sky, MT, USA, 2022; pp 01–12. 10.1109/AERO53065.2022.9843393.

(3) Sun, H.; Duan, M.; Wu, Y.; Zeng, Y.; Zhao, H.; Wu, S.; Lin, B.; Yang, R.; Tan, G. Designing Sustainable Built Environments for Mars Habitation: Integrating Innovations in Architecture, Systems, and Human Well-Being. Nexus 2024, 1 (3), 100030. 10.1016/j.ynexs.2024.100030.

(4) Pagnini, F.; Manzey, D.; Rosnet, E.; Ferravante, D.; White, O.; Smith, N. Human Behavior and Performance in Deep Space Exploration: Next Challenges and Research Gaps. npj Microgravity 2023, 9 (1), 27. 10.1038/s41526-023-00270-7.

(5) Koehle, A. P.; Brumwell, S. L.; Seto, E. P.; Lynch, A. M.; Urbaniak, C. Microbial Applications for Sustainable Space Exploration beyond Low Earth Orbit. npj Microgravity 2023, 9 (1), 47. 10.1038/s41526-023-00285-0.

(6) Weislogel, M. M.; Graf, J. C.; Wollman, A. P.; Turner, C. C.; Cardin, K. J. T.; Torres, L. J.; Goodman, J. E.; Buchli, J. C. How Advances in Low-g Plumbing Enable Space Exploration. npj Microgravity 2022, 8 (1), 16. 10.1038/s41526-022-00201-y.

(7) Silva, J. C. F.; Machado, K. L. D. G.; Silva, A. F. D. S.; Dias, R.; Bodnar, V. R.; Vieira, W. O.; Moreno-Pizani, M. A.; Ramos, J. D.; Santos, E. R. D.; Pauli, I.; Costa, L. C. Challenges and Opportunities for New Frontiers and Technologies to Guarantee Food Production: A Broad Systematic Perspective. January 15, 2025. 10.20944/preprints202501.1140.v1.

(8) Afshinnekoo, E.; Scott, R. T.; MacKay, M. J.; Pariset, E.; Cekanaviciute, E.; Barker, R.; Gilroy, S.; Hassane, D.; Smith, S. M.; Zwart, S. R.; Nelman-Gonzalez, M.; Crucian, B. E.; Ponomarev, S. A.; Orlov, O. I.; Shiba, D.; Muratani, M.; Yamamoto, M.; Richards, S. E.; Vaishampayan, P. A.; Meydan, C.; Foox, J.; Myrrhe, J.; Istasse, E.; Singh, N.; Venkateswaran, K.; Keune, J. A.; Ray, H. E.; Basner, M.; Miller, J.; Vitaterna, M. H.; Taylor, D. M.; Wallace, D.; Rubins, K.; Bailey, S. M.; Grabham, P.; Costes, S. V.; Mason, C. E.; Beheshti, A. Fundamental Biological Features of Spaceflight: Advancing the Field to Enable Deep-Space Exploration. Cell 2020, 183 (5), 1162–1184. 10.1016/j.cell.2020.10.050.

(9) Laranjeiro, R.; Harinath, G.; Pollard, A. K.; Gaffney, C. J.; Deane, C. S.; Vanapalli, S. A.; Etheridge, T.; Szewczyk, N. J.; Driscoll, M. Spaceflight Affects Neuronal Morphology and Alters Transcellular Degradation of Neuronal Debris in Adult Caenorhabditis Elegans. iScience 2021, 24 (2), 102105. 10.1016/j.isci.2021.102105.

(10) Lee, H.; Diao, J.; Tian, Y.; Guleria, R.; Lee, E.; Smith, A.; Savage, M.; Yeh, D.; Roberson, L.; Blenner, M.; Tang, Y. J.; Moon, T. S. Developing an Alternative Medium for In-Space Biomanufacturing. Nat Commun 2025, 16 (1), 728. 10.1038/s41467-025-56088-2.

(11) Fujimoto, M.; Tasker, E. J. Unveiling the Secrets of a Habitable World with JAXA’s Small-Body Missions. Nat Astron 2019, 3 (4), 284–286. 10.1038/s41550-019-0745-8.

(12) Creech, S.; Guidi, J.; Elburn, D. Artemis: An Overview of NASA’s Activities to Return Humans to the Moon. In 2022 IEEE Aerospace Conference (AERO); IEEE: Big Sky, MT, USA, 2022; pp 1–7. 10.1109/AERO53065.2022.9843277.

(13) Gu, Y. The China Space Station: A New Opportunity for Space Science. National Science Review 2022, 9 (1), nwab219. 10.1093/nsr/nwab219.

(14) Padma, T. V. India Shoots for the Moon with Chandrayaan-3 Lunar Lander. Nature 2023, d41586-023–02217–0. 10.1038/d41586-023-02217-0.

(15) Yin, Z.; Guo, P.; Wang, Z.; Yang, J.; Song, Y.; Zhou, Y.; Liu, M.; Cheng, C.; Wang, A. Review of the Development of Energy Technology Experiment for Space Station. Space Sci Technol 2026, 6, 0500. 10.34133/space.0500.

(16) Zhou, H.; Jiao, B.; Hou, Z.; Dang, Z. Satellite Formation Control for Cooperative Observation around the Rotating China Space Station. Space Sci Technol 2026, 6, 0348. 10.34133/space.0348.

(17) Wang, X.; Zhang, Q.; Wang, W. Design and Application Prospect of China’s Tiangong Space Station. Space Sci Technol 2023, 3, 0035. 10.34133/space.0035.

(18) Jia, Y.; Ma, Q.; Jian, P.; Liu, D.; Di, K.; Liu, B.; Mu, Y.; Xue, N.; Li, C.; Li, W.; Liu, W.; Liu, J.; Wan, G. A Survey on Path Planning Technologies for Lunar Long-Range Exploration Rovers. Space Sci Technol 2026, space.0646. 10.34133/space.0646.

(19) Youhanna, V.; Felicetti, L.; Ignatyev, D. Mars Planetary Insights and Design Framework for Future In-Situ Aerial Robotic Missions. Commun Eng 2026, 5 (1), 96. 10.1038/s44172-026-00647-y.

(20) Rosu, D.; Ceobanu, C. Space Education Activities To Encourage Stem Studies - A Theoretical Perspective; 2023; pp 626–632. 10.15405/epes.23056.58.

(21) Mardon, A. SPACE EXPLORATION: A Journey Through the Cosmos; De Gruyter, 2025. 10.1515/9781501521263.

(22) De Micco, V.; Amitrano, C.; Mastroleo, F.; Aronne, G.; Battistelli, A.; Carnero-Diaz, E.; De Pascale, S.; Detrell, G.; Dussap, C.-G.; Ganigué, R.; Jakobsen, Ø. M.; Poulet, L.; Van Houdt, R.; Verseux, C.; Vlaeminck, S. E.; Willaert, R.; Leys, N. Plant and Microbial Science and Technology as Cornerstones to Bioregenerative Life Support Systems in Space. npj Microgravity 2023, 9 (1), 69. 10.1038/s41526-023-00317-9.

(23) Zeidler, C.; Zabel, P.; Vrakking, V.; Dorn, M.; Bamsey, M.; Schubert, D.; Ceriello, A.; Fortezza, R.; De Simone, D.; Stanghellini, C.; Kempkes, F.; Meinen, E.; Mencarelli, A.; Swinkels, G.-J.; Paul, A.-L.; Ferl, R. J. The Plant Health Monitoring System of the EDEN ISS Space Greenhouse in Antarctica During the 2018 Experiment Phase. Front. Plant Sci. 2019, 10, 1457. 10.3389/fpls.2019.01457.

(24) Human Spaceflight and Exploration; Norberg, C., Ed.; Springer Berlin Heidelberg: Berlin, Heidelberg, 2013. 10.1007/978-3-642-23725-6.

(25) Space Age Science and Technology. Nature 1966, 209 (5022), 459–460. 10.1038/209459a0.

(26) Chavers, D. G.; Suzuki, M. N.; Smith, M. M.; Watson-Morgan, D. L.; Clarke, S. W.; Engelund, W. C.; Aitchison, M. L.; McEniry, M. S.; Means, L.; DeKlotz, M. M.; Jackson, S. IAF HUMAN SPACEFLIGHT SYMPOSIUM (B3) Governmental Human Spaceflight Programs (RFOverview) (1). th International Astronautical Congress 2019.

(27) Nowak, R. NASA’s Space Biology Program Shows Signs of Life. Science 1995, 268 (5210), 497–498. 10.1126/science.7725092.

(28) Fenn, W. O. The Challenge of Space Biology. AIBS Bulletin 1958, 8 (2), 15. 10.2307/1291925.

(29) Our Microbiology Correspondent. Space Biology: Regenerative Life Support. Nature 1969, 224 (5215), 112–112. 10.1038/224112a0.

(30) Stenzel, C. MATERIALS SCIENCE EXPERIMENTS UNDER MICROGRAVITY - A REVIEW OF HISTORY, FACILITIES, AND FUTURE OPPORTUNITIES; Marshall Space Flight Center.

(31) Monteiro, F.; Marques, P.; Simonini, A.; Carbonnelle, L.; Mendez, M. A. Experimental Characterization of Non-Isothermal Sloshing in Microgravity. arXiv 2024. 10.48550/ARXIV.2410.06590.

(32) Rezapour Sarabi, M.; Yetisen, A. K.; Tasoglu, S. Bioprinting in Microgravity. ACS Biomater. Sci. Eng. 2023, 9 (6), 3074–3083. 10.1021/acsbiomaterials.3c00195.

(33) Rutter, L. A.; Cope, H.; MacKay, M. J.; Herranz, R.; Das, S.; Ponomarev, S. A.; Costes, S. V.; Paul, A. M.; Barker, R.; Taylor, D. M.; Bezdan, D.; Szewczyk, N. J.; Muratani, M.; Mason, C. E.; Giacomello, S. Astronaut Omics and the Impact of Space on the Human Body at Scale. Nat Commun 2024, 15 (1), 4952. 10.1038/s41467-024-47237-0.

(34) Bushnell, D. M.; Moses, R. W. Prospectives in Deep Space Infrastructures, Development, and Colonization. NASA 2020.

(35) Lawrence, D. B.; Greco, J. A.; Birge, R. R.; Wagner, N. L. Layer-by-Layer Deposition in Microgravity for Enhanced Thin-Film Production. In In-Space Manufacturing and Resources; Hessel, V., Stoudemire, J., Miyamoto, H., Fisk, I. D., Eds.; Wiley, 2022; pp 285–301. 10.1002/9783527830909.ch15.

(36) Cockell, C. S.; Santomartino, R.; Finster, K.; Waajen, A. C.; Eades, L. J.; Moeller, R.; Rettberg, P.; Fuchs, F. M.; Van Houdt, R.; Leys, N.; Coninx, I.; Hatton, J.; Parmitano, L.; Krause, J.; Koehler, A.; Caplin, N.; Zuijderduijn, L.; Mariani, A.; Pellari, S. S.; Carubia, F.; Luciani, G.; Balsamo, M.; Zolesi, V.; Nicholson, N.; Loudon, C.-M.; Doswald-Winkler, J.; Herová, M.; Rattenbacher, B.; Wadsworth, J.; Craig Everroad, R.; Demets, R. Space Station Biomining Experiment Demonstrates Rare Earth Element Extraction in Microgravity and Mars Gravity. Nat Commun 2020, 11 (1), 5523. 10.1038/s41467-020-19276-w.

(37) Barker, R.; Kruse, C. P. S.; Johnson, C.; Saravia-Butler, A.; Fogle, H.; Chang, H.-S.; Trane, R. M.; Kinscherf, N.; Villacampa, A.; Manzano, A.; Herranz, R.; Davin, L. B.; Lewis, N. G.; Perera, I.; Wolverton, C.; Gupta, P.; Jaiswal, P.; Reinsch, S. S.; Wyatt, S.; Gilroy, S. Meta-Analysis of the Space Flight and Microgravity Response of the Arabidopsis Plant Transcriptome. npj Microgravity 2023, 9 (1), 21. 10.1038/s41526-023-00247-6

(38) Ferl, R.; Wheeler, R.; Levine, H. G.; Paul, A.-L. Plants in Space. Current Opinion in Plant Biology 2002, 5 (3), 258–263. 10.1016/S1369-5266(02)00254-6.

(39) Ohnishi, T. Life Science Experiments Performed in Space in the ISS/Kibo Facility and Future Research Plans. J Radiat Res 2016, 57 (S1), i41–i46. 10.1093/jrr/rrw020.

(40) Overbey, E. G.; Saravia-Butler, A. M.; Zhang, Z.; Rathi, K. S.; Fogle, H.; Da Silveira, W. A.; Barker, R. J.; Bass, J. J.; Beheshti, A.; Berrios, D. C.; Blaber, E. A.; Cekanaviciute, E.; Costa, H. A.; Davin, L. B.; Fisch, K. M.; Gebre, S. G.; Geniza, M.; Gilbert, R.; Gilroy, S.; Hardiman, G.; Herranz, R.; Kidane, Y. H.; Kruse, C. P. S.; Lee, M. D.; Liefeld, T.; Lewis, N. G.; McDonald, J. T.; Meller, R.; Mishra, T.; Perera, I. Y.; Ray, S.; Reinsch, S. S.; Rosenthal, S. B.; Strong, M.; Szewczyk, N. J.; Tahimic, C. G. T.; Taylor, D. M.; Vandenbrink, J. P.; Villacampa, A.; Weging, S.; Wolverton, C.; Wyatt, S. E.; Zea, L.; Costes, S. V.; Galazka, J. M. NASA GeneLab RNA-Seq Consensus Pipeline: Standardized Processing of Short-Read RNA-Seq Data. iScience 2021, 24 (4), 102361. 10.1016/j.isci.2021.102361.

(41) Fong, K. Moon Landing: Space Medicine and the Legacy of Project Apollo. The Lancet 2019, 394 (10194), 205–207. 10.1016/S0140-6736(19)31568-5.

(42) Overbey, E. G.; Kim, J.; Tierney, B. T.; Park, J.; Houerbi, N.; Lucaci, A. G.; Garcia Medina, S.; Damle, N.; Najjar, D.; Grigorev, K.; Afshin, E. E.; Ryon, K. A.; Sienkiewicz, K.; Patras, L.; Klotz, R.; Ortiz, V.; MacKay, M.; Schweickart, A.; Chin, C. R.; Sierra, M. A.; Valenzuela, M. F.; Dantas, E.; Nelson, T. M.; Cekanaviciute, E.; Deards, G.; Foox, J.; Narayanan, S. A.; Schmidt, C. M.; Schmidt, M. A.; Schmidt, J. C.; Mullane, S.; Tigchelaar, S. S.; Levitte, S.; Westover, C.; Bhattacharya, C.; Lucotti, S.; Wain Hirschberg, J.; Proszynski, J.; Burke, M.; Kleinman, A. S.; Butler, D. J.; Loy, C.; Mzava, O.; Lenz, J.; Paul, D.; Mozsary, C.; Sanders, L. M.; Taylor, L. E.; Patel, C. O.; Khan, S. A.; Suhail Mohamad, M.; Byhaqui, S. G. A.; Aslam, B.; Gajadhar, A. S.; Williamson, L.; Tandel, P.; Yang, Q.; Chu, J.; Benz, R. W.; Siddiqui, A.; Hornburg, D.; Blease, K.; Moreno, J.; Boddicker, A.; Zhao, J.; Lajoie, B.; Scott, R. T.; Gilbert, R. R.; Lai Polo, S.; Altomare, A.; Kruglyak, S.; Levy, S.; Ariyapala, I.; Beer, J.; Zhang, B.; Hudson, B. M.; Rininger, A.; Church, S. E.; Beheshti, A.; Church, G. M.; Smith, S. M.; Crucian, B. E.; Zwart, S. R.; Matei, I.; Lyden, D. C.; Garrett-Bakelman, F.; Krumsiek, J.; Chen, Q.; Miller, D.; Shuga, J.; Williams, S.; Nemec, C.; Trudel, G.; Pelchat, M.; Laneuville, O.; De Vlaminck, I.; Gross, S.; Bolton, K. L.; Bailey, S. M.; Granstein, R.; Furman, D.; Melnick, A. M.; Costes, S. V.; Shirah, B.; Yu, M.; Menon, A. S.; Mateus, J.; Meydan, C.; Mason, C. E. The Space Omics and Medical Atlas (SOMA) and International Astronaut Biobank. Nature 2024, 632 (8027), 1145–1154. 10.1038/s41586-024-07639-y.

(43) Sanders, L. M.; Lopez, D. K.; Wood, A. E.; Scott, R. T.; Gebre, S. G.; Saravia-Butler, A. M.; Costes, S. V. Celebrating 30 Years of Access to NASA Space Life Sciences Data. GigaScience 2024, 13, giae066. 10.1093/gigascience/giae066.

(44) Stoehr, F.; Manning, A.; McLay, S.; Ashigatawa, K.; del Prado, M.; Jenkins, D.; Damian, A.; Wang, K.-S.; Moraghan, A.; Plunkett, A.; Lipnicky, A.; Sanhueza, P.; Rivera, G. C.; Gaudet, S. The ALMA Science Archive Reaches a Major Milestone. 2022. 10.48550/ARXIV.2208.03275.

(45) Blumberg, B. S. The NASA Astrobiology Institute: Early History and Organization. Astrobiology 2003, 3 (3), 463–470. 10.1089/153110703322610573.

(46) Rudenko, M. Minor Planet Center: Data Processing Challenges. Proc. IAU 2015, 10 (S318), 265–269. 10.1017/S174392131500839X.

(47) Lin, Y.; Yang, W.; Zhang, H.; Hui, H.; Hu, S.; Xiao, L.; Liu, J.; Xiao, Z.; Yue, Z.; Zhang, J.; Liu, Y.; Yang, J.; Lin, H.; Zhang, A.; Guo, D.; Gou, S.; Xu, L.; He, Y.; Zhang, X.; Qin, L.; Ling, Z.; Li, X.; Du, A.; He, H.; Zhang, P.; Cao, J.; Li, X. Return to the Moon: New Perspectives on Lunar Exploration. Science Bulletin 2024, 69 (13), 2136–2148. 10.1016/j.scib.2024.04.051.

(48) García, R. F. MVC: Model–View–Controller. In iOS Architecture Patterns; Apress: Berkeley, CA, 2023; pp 45–106. 10.1007/978-1-4842-9069-9_2.

(49) Thakur, P.; Jadon, P. Django: Developing Web Using Python. In 2023 3rd International Conference on Advance Computing and Innovative Technologies in Engineering (ICACITE); IEEE: Greater Noida, India, 2023; pp 303–306. 10.1109/ICACITE57410.2023.10183246.

(50) Salunke, S. V.; Ouda, A. A Performance Benchmark for the PostgreSQL and MySQL Databases. Future Internet 2024, 16 (10), 382. 10.3390/fi16100382.

(51) Milutinović, V.; Salom, J.; Trifunovic, N.; Giorgi, R. Using the WebIDE. In Guide to DataFlow Supercomputing; Computer Communications and Networks; Springer International Publishing: Cham, 2015; pp 107–122. 10.1007/978-3-319-16229-4_4.

(52) Benson, D. A.; Cavanaugh, M.; Clark, K.; Karsch-Mizrachi, I.; Lipman, D. J.; Ostell, J.; Sayers, E. W. GenBank. Nucleic Acids Research 2012, 41 (D1), D36–D42. 10.1093/nar/gks1195.

(53) Fabien, C. Stellarium. Stellarium.

(54) Reimers, N.; Gurevych, I. Making Monolingual Sentence Embeddings Multilingual Using Knowledge Distillation. arXiv 2020. 10.48550/ARXIV.2004.09813.

(55) Beltagy, I.; Lo, K.; Cohan, A. SciBERT: A Pretrained Language Model for Scientific Text. In Proceedings of the 2019 Conference on Empirical Methods in Natural Language Processing and the 9th International Joint Conference on Natural Language Processing (EMNLP-IJCNLP); Association for Computational Linguistics: Hong Kong, China, 2019; pp 3613–3618. 10.18653/v1/D19-1371.

(56) Deka, P.; Jurek-Loughrey, A.; Deepak. Unsupervised Keyword Combination Query Generation from Online Health Related Content for Evidence-Based Fact Checking. In The 23rd International Conference on Information Integration and Web Intelligence; ACM: Linz Austria, 2021; pp 267–277. 10.1145/3487664.3487701.

(57) Rousseeuw, P. J. Silhouettes: A Graphical Aid to the Interpretation and Validation of Cluster Analysis. Journal of Computational and Applied Mathematics 1987, 20, 53–65. 10.1016/0377-0427(87)90125-7.

(58) Xiao, J.; Lu, J.; Li, X. Davies Bouldin Index Based Hierarchical Initialization K-Means. IDA 2017, 21 (6), 1327–1338. 10.3233/IDA-163129.

(59) Lukasik, S.; Kowalski, P. A.; Charytanowicz, M.; Kulczycki, P. Clustering Using Flower Pollination Algorithm and Calinski-Harabasz Index. In 2016 IEEE Congress on Evolutionary Computation (CEC); IEEE: Vancouver, BC, Canada, 2016; pp 2724–2728. 10.1109/CEC.2016.7744132.

(60) Likas, A.; Vlassis, N.; J. Verbeek, J. The Global K-Means Clustering Algorithm. Pattern Recognition 2003, 36 (2), 451–461. 10.1016/S0031-3203(02)00060-2.

(61) Hricik, T.; Bader, D.; Green, O. Using RAPIDS AI to Accelerate Graph Data Science Workflows. In 2020 IEEE High Performance Extreme Computing Conference (HPEC); IEEE: Waltham, MA, USA, 2020; pp 1–4. 10.1109/HPEC43674.2020.9286224.

(62) Hricik, T.; Bader, D.; Green, O. Using RAPIDS AI to Accelerate Graph Data Science Workflows. In 2020 IEEE High Performance Extreme Computing Conference (HPEC); IEEE: Waltham, MA, USA, 2020; pp 1–4. 10.1109/HPEC43674.2020.9286224.

(63) McInnes, L.; Healy, J.; Melville, J. UMAP: Uniform Manifold Approximation and Projection for Dimension Reduction. arXiv 2018. 10.48550/ARXIV.1802.03426.

(64) Abu Alfeilat, H. A.; Hassanat, A. B. A.; Lasassmeh, O.; Tarawneh, A. S.; Alhasanat, M. B.; Eyal Salman, H. S.; Prasath, V. B. S. Effects of Distance Measure Choice on K-Nearest Neighbor Classifier Performance: A Review. Big Data 2019, 7 (4), 221–248. 10.1089/big.2018.0175.

(65) Overbey, E. G.; Kim, J.; Tierney, B. T.; Park, J.; Houerbi, N.; Lucaci, A. G.; Garcia Medina, S.; Damle, N.; Najjar, D.; Grigorev, K.; Afshin, E. E.; Ryon, K. A.; Sienkiewicz, K.; Patras, L.; Klotz, R.; Ortiz, V.; MacKay, M.; Schweickart, A.; Chin, C. R.; Sierra, M. A.; Valenzuela, M. F.; Dantas, E.; Nelson, T. M.; Cekanaviciute, E.; Deards, G.; Foox, J.; Narayanan, S. A.; Schmidt, C. M.; Schmidt, M. A.; Schmidt, J. C.; Mullane, S.; Tigchelaar, S. S.; Levitte, S.; Westover, C.; Bhattacharya, C.; Lucotti, S.; Wain Hirschberg, J.; Proszynski, J.; Burke, M.; Kleinman, A. S.; Butler, D. J.; Loy, C.; Mzava, O.; Lenz, J.; Paul, D.; Mozsary, C.; Sanders, L. M.; Taylor, L. E.; Patel, C. O.; Khan, S. A.; Suhail Mohamad, M.; Byhaqui, S. G. A.; Aslam, B.; Gajadhar, A. S.; Williamson, L.; Tandel, P.; Yang, Q.; Chu, J.; Benz, R. W.; Siddiqui, A.; Hornburg, D.; Blease, K.; Moreno, J.; Boddicker, A.; Zhao, J.; Lajoie, B.; Scott, R. T.; Gilbert, R. R.; Lai Polo, S.; Altomare, A.; Kruglyak, S.; Levy, S.; Ariyapala, I.; Beer, J.; Zhang, B.; Hudson, B. M.; Rininger, A.; Church, S. E.; Beheshti, A.; Church, G. M.; Smith, S. M.; Crucian, B. E.; Zwart, S. R.; Matei, I.; Lyden, D. C.; Garrett-Bakelman, F.; Krumsiek, J.; Chen, Q.; Miller, D.; Shuga, J.; Williams, S.; Nemec, C.; Trudel, G.; Pelchat, M.; Laneuville, O.; De Vlaminck, I.; Gross, S.; Bolton, K. L.; Bailey, S. M.; Granstein, R.; Furman, D.; Melnick, A. M.; Costes, S. V.; Shirah, B.; Yu, M.; Menon, A. S.; Mateus, J.; Meydan, C.; Mason, C. E. The Space Omics and Medical Atlas (SOMA) and International Astronaut Biobank. Nature 2024, 632 (8027), 1145–1154. 10.1038/s41586-024-07639-y.

(66) Al-Ahmadi, W.; Barnawi, R.; Hitti, E. G.; Khabar, K. S. A. Spaceflight Alters Molecular Networks Linked to Diverse Human Diseases in a Single Cellular Model. Sci. Adv. 2026, 12 (1), eadw7832. 10.1126/sciadv.adw7832.

(67) Wadhwa, A.; Moreno-Villanueva, M.; Crucian, B.; Wu, H. Synergistic Interplay between Radiation and Microgravity in Spaceflight-Related Immunological Health Risks. Immun Ageing 2024, 21 (1), 50. 10.1186/s12979-024-00449-w.

(68) Giacinto, O.; Lusini, M.; Sammartini, E.; Minati, A.; Mastroianni, C.; Nenna, A.; Pascarella, G.; Sammartini, D.; Carassiti, M.; Miraldi, F.; Chello, M.; Pelliccia, F. Cardiovascular Effects of Cosmic Radiation and Microgravity. JCM 2024, 13 (2), 520. 10.3390/jcm13020520.

(69) Witze, A. Lift off! Artemis II Mission Sends Humans to the Moon — Opening a New Era of Exploration. Nature 2026, 652 (8109), 279–281. 10.1038/d41586-026-00978-y.

(70) Covey, C. Moon and Venus as Worthy of Exploration as Mars. Nature 2007, 445 (7125), 256–256. 10.1038/445256c.

(71) Chatzitheodoridis, E.; De Vera, J.-P.; Kereszturi, A.; Mason, N.; Possnig, C.; Puumala, M.; Sivula, O.; Viso, M.; Arnould, J.; Detrell, G.; Ewald, R.; Freissinet, C.; Holynska, M.; Javaux, E.; Lee, N. M.; Lehto, K.; Milligan, T.; Siljeström, S.; Vago, J. L. Perspectives for Crewed Missions to Mars: Exploration from Orbit and/or Short Stay. In Mars and the Earthlings: A Realistic View on Mars Exploration and Settlement; Verseux, C., Gargaud, M., Lehto, K., Viso, M., Eds.; Space and Society; Springer Nature Switzerland: Cham, 2024; pp 117–197. 10.1007/978-3-031-66881-4_6.

(72) Markley, R. Moving beyond “Why Mars?” For the Love of Mars: A Human History of the Red Planet Matthew Shindell University of Chicago Press, 2023. 248 Pp. Science 2023, 380 (6646), 697–697. 10.1126/science.adh7737.

(73) NASA Lays out Vision for Robotic Mars Exploration, 2023. 10.1126/science.adi0331.

(74) Gebre, S. G.; Scott, R. T.; Saravia-Butler, A. M.; Lopez, D. K.; Sanders, L. M.; Costes, S. V. NASA Open Science Data Repository: Open Science for Life in Space. Nucleic Acids Research 2025, 53 (D1), D1697–D1710. 10.1093/nar/gkae1116.

